# A rapid, machine-learning readout of cochlear neural degeneration that transfers from animals to humans

**DOI:** 10.64898/2026.09.24.754199

**Authors:** Luz S. Andrino, Richard Gall, Jennifer Klara, Murat Akcakaya, Edward L. Bartlett, Aravindakshan Parthasarathy

**Author notes:** **Corresponding author:** Aravindakshan Parthasarathy, Department of Communication Science and Disorders, University of Pittsburgh, Pittsburgh, PA 15213.

## Abstract

Standard hearing tests miss cochlear neural degeneration (CND), the loss of synapses between inner hair cells and auditory nerve fibers that may underlie unexplained hearing difficulty in roughly one in ten adults. Envelope following responses (EFRs) can index this neural loss noninvasively, but conventional protocols are too slow for clinical use. We tested a dynamic amplitude-modulated (dAM) EFR that sweeps the full modulation spectrum in a single brief stimulus. In Mongolian gerbils with histologically verified CND, dAM EFRs revealed selective deficits at fast modulation rates without threshold elevation. A machine-learning classifier distinguished young from middleaged animals with high accuracy; additionally, its most informative feature, power near 400-500 Hz, tracked synapse counts, indicating a readout of cochlear neural health rather than age. Applied without retraining to 56 human listeners, the gerbil-trained classifier separated age groups above chance. dAM EFRs thus offer a rapid, objective and clinically viable marker of CND.

## Introduction

Difficulty understanding speech-in-noise is the most common complaint among adults seeking hearing healthcare, yet clinical practice still centers on pure-tone audiometry and the restoration of audibility in quiet. As a result, approximately 1 in 10 adults who present to hearing clinics report substantial hearing difficulties despite audiometric thresholds within normal limits (*1*–*4*). These patients lack an objective biomarker for their complaint, have no established intervention, and have little diagnostic basis to justify accommodations, leaving a large subpopulation effectively invisible to current diagnostic tools and underserved by current care. Closing this gap requires a clinically deployable test that detects pathologies underlying suprathreshold listening difficulty in the absence of threshold elevations.

A growing body of evidence implicates cochlear neural degeneration (CND) as the leading candidate mechanism underlying these hidden deficits. CND is defined by the loss of afferent connections between the cochlea and brain, including degeneration of inner hair cell-auditory nerve synapses, spiral ganglion neurons, or their myelination (*5*–*9*). Across every mammalian species studied so far, including non-human primates, aging and noise exposure produce a substantial loss of cochlear afferent synapses often exceeding 50%, well before any elevation of audiometric threshold (*10*–*14*). Human temporal-bone histology provides further evidence of extensive age- and noise-related afferent neural loss, with deafferentation reaching approximately 40% by the fifth decade of life (*15*–*17*). Because hearing thresholds do not reflect information lost at degraded neural synapses, CND offers a convincing explanation for why listeners with normal audiograms experience suprathreshold difficulty.

Envelope following responses (EFRs) are a promising non-invasive candidate biomarker for cochlear neural degeneration, combining suprathreshold sensitivity, anatomical specificity, and a direct link to histology (*18*–*24*). EFRs are scalp-recorded, neural responses that are phase-locked to the temporal envelope of amplitude-modulated sounds. Because the upper limit of neural phase-locking for amplitude modulation decreases systematically from the periphery to the cortex, varying the modulation rate shifts the dominant EFR generator along the ascending pathway (*25*). Slow rates (below ∼80 Hz) emphasize cortical sources (*26, 27*), whereas fast rates (above ∼400 Hz) are dominated by the auditory nerve and cochlear nucleus,(*18, 19, 28*) the structure most directly affected by CND. In rodent models, suprathreshold EFR amplitudes at fast modulation rates correlate with histologically counted afferent synapses and decline with age and noise exposure before threshold elevation (*18, 19, 29, 30*). Middle-aged adults who, despite normal audiograms, report increased listening effort and speech-in-noise difficulty also exhibit near-identical age-related EFR declines at fast modulation rates to those observed in rodent models(*28, 29*), establishing high-modulation-rate EFRs as a sensitive, mechanistically grounded marker of age-related CND.

Despite this mechanistic suitability, conventional envelope following response protocols face practical barriers to clinical use. EFRs are typically recorded one modulation frequency at a time, and characterizing the temporal modulation transfer function across the physiologically relevant range requires testing times that are unfeasible in the clinic. Furthermore, the substantial time demands of these lengthy protocols can limit research studies to sampling often just one or a small number of modulation rates. This risks omission of diagnostic features most informative for a given individual, whose neural responses with specific modulation rates may vary with head size, neural-generator geometry, and recording conditions. Therefore, any clinically viable measure will require reduced acquisition time and increased spectrotemporal resolution while preserving the high modulation rate sensitivity that makes EFRs informative about cochlear neural degeneration.

We recently developed a dynamic amplitude-modulated (dAM) EFR paradigm that addresses these barriers (*31*). A single continuously sweeping stimulus traverses the full modulation range (16-1500 Hz) in 1 second, and a spectrally specific frequency demodulation analysis aligns the time-varying response to its known trajectory, circumventing the trade-off between the length of the analysis window and spectral resolution (*32*). This yields an approximately 30-fold improvement in spectrotemporal resolution and a roughly 5-fold reduction in recording time relative to the conventional sinusoidally amplitude-modulated (sAM) protocols, with responses that correlate strongly with routine EFR metrics. dAM EFRs have been validated across four mammalian species, including humans. Here, we tested if dAM EFRs are sensitive to cochlear neural degeneration, by recording discrete sAM EFRs, and dAM EFRs in young and middle-aged Mongolian gerbils with histologically verified cochlear synapse counts. We show that (1) middleaged gerbils exhibit cochlear synapse loss with preserved auditory thresholds and selective EFR reductions at fast modulation rates with both paradigms and (2) machine-learning classification of dAM EFR spectral features discriminates age groups with high accuracy, with the most discriminative feature, power in the 400-500 Hz band, correlating directly with synapse counts. To test the human translational feasibility of the dAM-classification framework to human subjects, we successfully applied the classifier trained on gerbils to dAM EFR responses recorded from middle-aged human listeners with clinically normal audiograms and putative CND. Together, these results validate dAM EFRs as a rapid, objective marker of age-related temporal-processing deficits and a candidate biomarker of CND primed for clinical translation.

## Results

### Middle-aged gerbils exhibit cochlear synapse loss without threshold elevation and selective EFR deficits at fast modulation rates

We studied young (28.3 ± 2.4 weeks; *n* = 24) and middle-aged (82.8 ± 1.5 weeks; *n* = 19) Mongolian gerbils (*Meriones unguiculatus*, Fig. 1A), a model whose low-frequency hearing range closely matches that of humans (*33*). Hearing thresholds measured using ABRs showed minimal threshold loss in middle-aged animals across tones from 0.5 to 8 kHz or to a broadband chirp, which while statistically significant, were all below 10 dB SPL, and considered clinically normal (Fig. 1B, Table 1). However, immunohistological analysis of cochlear whole-mounts revealed significant age-related loss of afferent ribbon synapses at multiple cochlear regions (Fig. 1C-D, Table 3,4), including at the 3 kHz region, which corresponds to the recording carrier frequency, where middle-aged animals showed an approximately 15% reduction in ribbons per inner hair cell (*t* = 3.00, *p* = 0.006, Cohen’s *d* = 0.97; Fig. 1D, inset). Together, these findings confirm age-related cochlear synaptic loss in the presence of clinically normal audiometric thresholds in our middleaged gerbil model.

**Table 1.** Two-way mixed ANOVA: ABR thresholds (Age Group × Frequency).

| Source | SS | df1 | df2 | MS | F | p | p (GG) | $\eta_p^2$ | $\epsilon$ |
| --- | --- | --- | --- | --- | --- | --- | --- | --- | --- |
| Age Group | 580.671 | 1 | 29 | 580.671 | 7.139 | .012 | — | .198 | — |
| Frequency | 5797.419 | 4 | 116 | 1449.355 | 24.880 | <.001 | <.001 | .462 | .749 |
| Interaction | 295.256 | 4 | 116 | 73.814 | 1.267 | .287 | — | .042 | — |

**Table 2.**
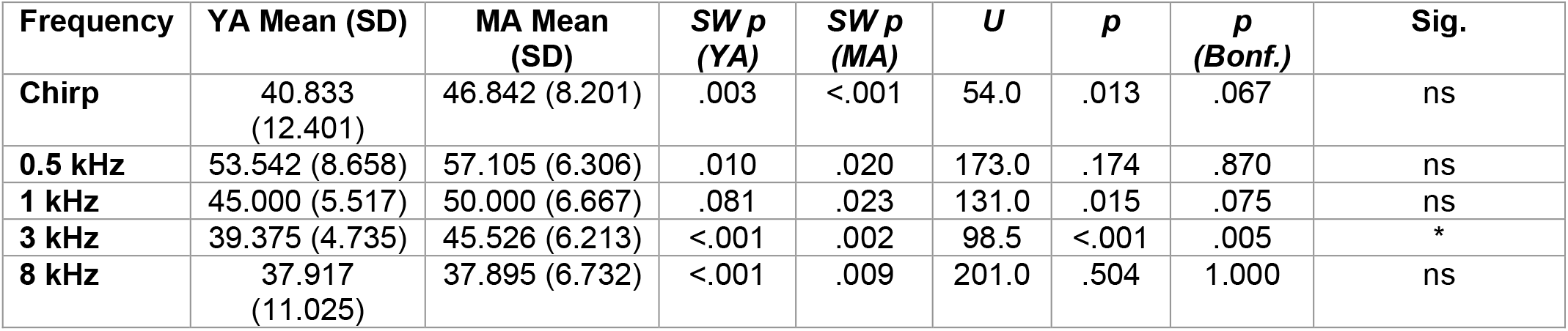
Shapiro–Wilk and Mann–Whitney U: ABR thresholds at each frequency.

**Table 3.** Two-way mixed ANOVA: synapse counts (Age Group × Cochlear Frequency Region).

| Source | SS | df1 | df2 | MS | F | p | p (GG) | $\eta_p^2$ | $\epsilon$ |
| --- | --- | --- | --- | --- | --- | --- | --- | --- | --- |
| Age Group | 191.146 | 1 | 18 | 191.146 | 23.330 | <.001 | — | .564 | — |
| Cochlear Freq. | 125.477 | 3 | 54 | 41.826 | 6.448 | <.001 | <.001 | .264 | .923 |
| Interaction | 2.888 | 3 | 54 | 0.963 | 0.148 | .930 | — | .008 | — |

**Table 4.** Shapiro–Wilk and Mann–Whitney U: Synapse Counts.

| Cochlear Region (kHz) | YA Mean (SD) | MA Mean (SD) | SW p (YA) | SW p (MA) | U | p | p (Bonf.) | Sig. |
| --- | --- | --- | --- | --- | --- | --- | --- | --- |
| 0.5 | 19.893 (3.292) | 17.020 (1.136) | .399 | .071 | 108.0 | .011 | .043 | * |
| 1 | 20.779 (4.067) | 19.934 (2.540) | .232 | .079 | 121.0 | .740 | 1.000 | ns |
| 3 | 23.365 (2.764) | 20.438 (2.576) | .858 | .184 | 175.5 | .010 | .038 | * |
| 8 | 20.930 (2.886) | 17.981 (1.821) | .720 | .028 | 124.0 | .011 | .044 | * |

**Figure 1.**
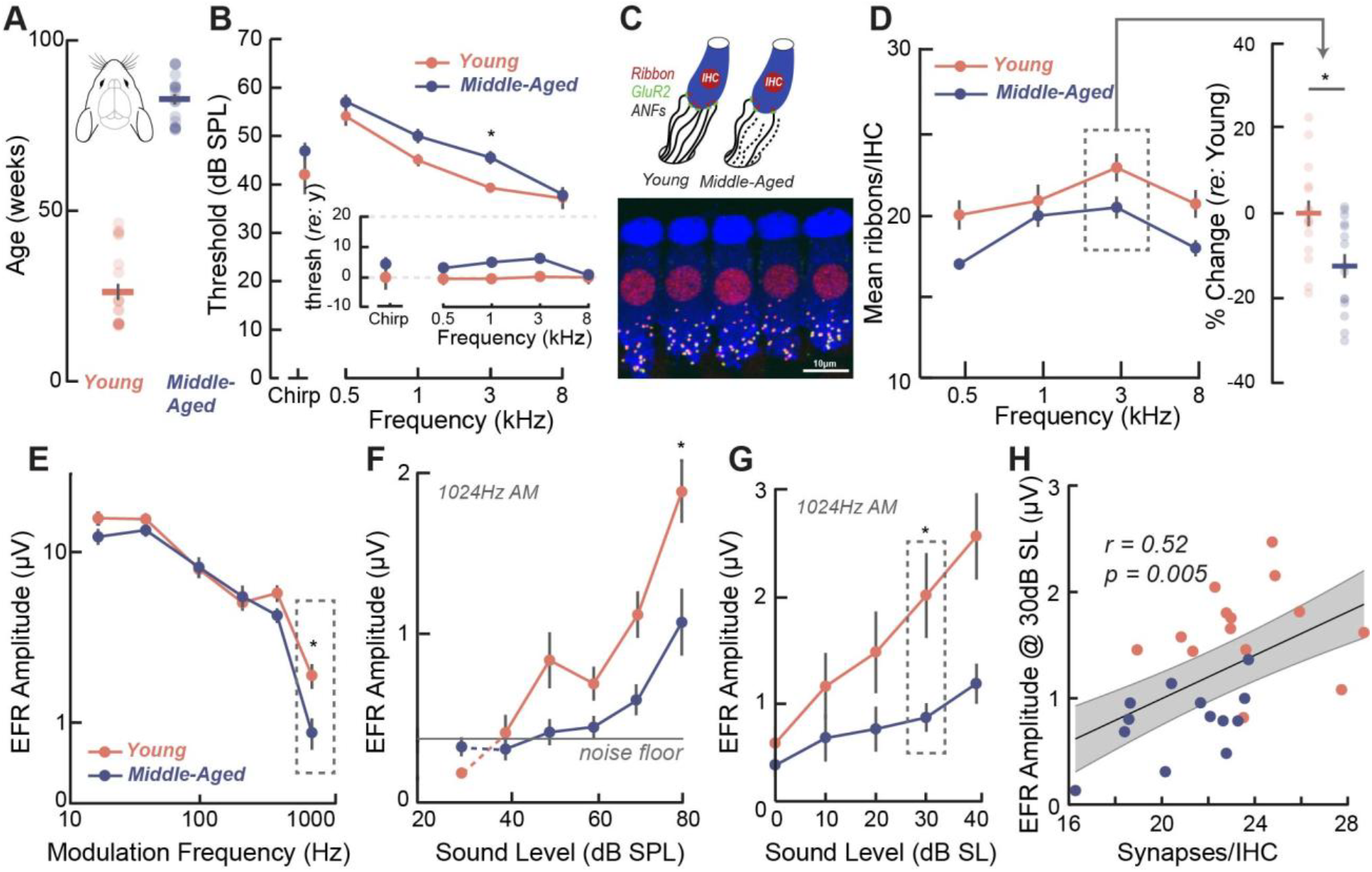
EFRs at fast modulation rates are associated with histologically confirmed age-related CND. **(A)** Age distribution of young (28.3±2.4, weeks, *n* = 24) and middle-aged (82.8±1.5 weeks, *n* = 19) gerbils. **(B)** Audiometric thresholds across frequency show minimal elevation in middle-aged relative to young animals (inset: thresholds relative to young), all within 10 dB SPL. **(C)** Schematic of the inner hair cell synaptic junction: presynaptic ribbons (CtBP2, red), postsynaptic glutamate receptor patches (GluR2, green), and auditory nerve fibers (ANFs, black) undergoing deafferentation with age (top); representative confocal image of hair cells in an immunostained cochlear whole mount (bottom). **(D)** Mean ribbon-synapse counts per inner hair cell across cochlear frequency regions (0.5-8 kHz) showing reduced ribbon synapse counts in middle-aged animals, particularly at the 3 kHz region (dashed box) corresponding to the EFR carrier frequency; inset shows percent change relative to young at the 3 kHz region. **(E)** EFR amplitude versus modulation frequency (16-1024 Hz) at 30 dB SL for young and middle-aged gerbils with selective reduction in middle-aged animals at fast rates (dashed box). **(F)** EFR amplitude at 1024 Hz across sound levels (20-80 dB SPL); middle-aged animals show reduced EFR amplitudes with increasing sound level. **(G)** EFR amplitude versus modulation frequency at equalized sensation level, correcting for individual hearing sensitivity further showed significant decreases in EFR amplitudes with age at suprathreshold sound levels **(H)** EFR amplitude at 1024 Hz (3 kHz carrier, 30 dB SL) correlated with synapses per inner hair cell at the 3 kHz region. Error bars = standard error of the mean (SEM). Asterisks = p < 0.05.

To assess the peripheral neural coding consequences of CND, we recorded EFRs to sinusoidally amplitude-modulated tones at a 3 kHz carrier across modulation frequencies from 16 to 1024 Hz. At a fixed individualized suprathreshold sensation level (30 dB SL) that accounts for the small variations in hearing thresholds with age, middle-aged gerbils showed EFR amplitudes comparable to young animals at slow modulation rates but reduced amplitudes at faster rates above 500 Hz with statistically significant reductions at 1024 Hz (Fig. 1E, Tables 5, 6), consistent with CND (*18, 19, 23, 29*). EFR growth functions at 1024 Hz across 20-80 dB SPL showed a significant age × level interaction, with the largest group difference at 80 dB SPL (Fig. 1F, Tables 7, 8). EFR amplitudes at 1024Hz AM replotted relative to individual animal thresholds further revealed suprathreshold EFR deficits at matched sensation levels with a significant main effect of age and a significant age × sensation level interaction with the largest group differences at 30dB SL (Fig. 1G, Tables 9, 10). Finally, EFR amplitudes at 1024 Hz and 30 dB SL significantly correlated with synapse counts at the 3 kHz region of the cochlea across animals, linking the physiological deficit to the underlying synaptopathy (Fig. 1H).

**Table 5.**
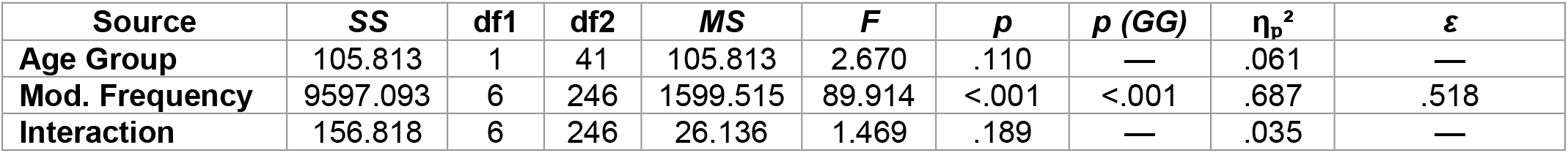
Two-way mixed ANOVA: sAM EFR amplitude (Age Group × Modulation Frequency).

**Table 6.**
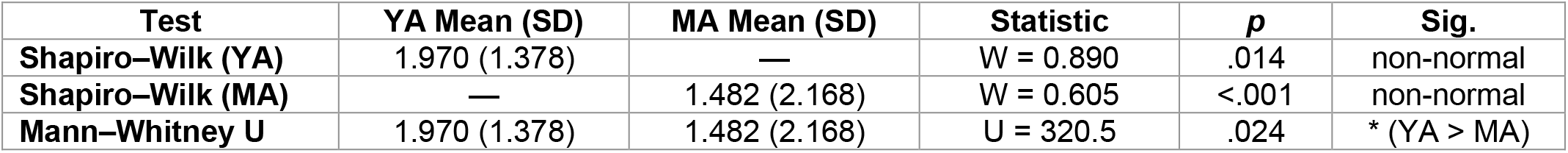
Shapiro–Wilk and Mann–Whitney U: sAM EFR amplitude at 1024 Hz AM (30 dB SL).

**Table 7.** Two-way mixed ANOVA: EFR amplitude (Age Group × Sound Level).

| Source | SS | df1 | df2 | MS | F | p | p (GG) | $\eta_p^2$ | $\varepsilon$ |
| --- | --- | --- | --- | --- | --- | --- | --- | --- | --- |
| Age Group | 9.412 | 1 | 41 | 9.412 | 3.780 | .059 | — | .084 | — |
| Sound Level | 84.727 | 6 | 246 | 14.121 | 33.404 | <.001 | <.001 | .449 | .375 |
| Interaction | 11.455 | 6 | 246 | 1.909 | 4.516 | <.001 | — | .099 | — |

**Table 8.** Mann–Whitney U: EFR amplitude at each sound level (20–80 dB SPL).

| Level (dB SPL) | YA Mean (SD) | MA Mean (SD) | U | p | p (Bonf.) | Sig. |
| --- | --- | --- | --- | --- | --- | --- |
| 20 | 0.154 (0.093) | 0.307 (0.321) | 212.0 | .705 | 1.000 | ns |
| 30 | 0.289 (0.271) | 0.260 (0.249) | 259.0 | .456 | 1.000 | ns |
| 40 | 0.508 (0.604) | 0.439 (0.527) | 223.0 | .912 | 1.000 | ns |
| 50 | 0.908 (0.755) | 0.533 (0.548) | 300.0 | .080 | .563 | ns |
| 60 | 1.147 (0.997) | 0.773 (1.044) | 310.0 | .046 | .324 | ns |
| 70 | 1.700 (1.380) | 0.947 (0.962) | 330.0 | .013 | .091 | ns |
| 80 | 2.213 (1.500) | 1.169 (0.902) | 340.0 | .006 | .045 | * |

**Table 9.** Two-way mixed ANOVA: EFR amplitude (Age Group × Sensation Level).

| Source | SS | df1 | df2 | MS | F | p | p (GG) | $\eta_p^2$ | $\varepsilon$ |
| --- | --- | --- | --- | --- | --- | --- | --- | --- | --- |
| Age Group | 11.188 | 1 | 34 | 11.188 | 10.556 | .003 | — | .237 | — |
| Sensation Level | 33.449 | 4 | 136 | 8.362 | 32.292 | <.001 | <.001 | .487 | .604 |
| Interaction | 2.908 | 4 | 136 | 0.727 | 2.808 | .028 | — | .076 | — |

**Table 10.** Mann–Whitney U: EFR amplitude at each sensation level.

| Level (dB SL) | YA Mean (SD) | MA Mean (SD) | U | p | p (Bonf.) | Sig. |
| --- | --- | --- | --- | --- | --- | --- |
| 0 | 0.482 (0.507) | 0.306 (0.203) | 189.0 | .898 | 1.000 | ns |
| 10 | 0.953 (0.713) | 0.553 (0.396) | 289.0 | .065 | .458 | ns |
| 20 | 1.091 (0.842) | 0.459 (0.265) | 293.0 | .008 | .056 | ns |
| 30 | 1.615 (0.971) | 0.733 (0.319) | 298.0 | .001 | .008 | * |
| 40 | 2.071 (1.112) | 1.382 (0.820) | 287.0 | .043 | .299 | ns |

### dAM EFRs detect the same CND-related deficits with greater efficiency

Discrete sAM EFRs require serial recording at each modulation rate which limits spectrotemporal resolution and sensitivity to modulation rate-specific deficits. To overcome these constraints, we recorded our recently developed dAM EFRs in the same animals using a single stimulus whose modulation frequency swept continuously from low to high rates (Figs. 2A-B). Representative responses, visualized as spectrograms, tracked the stimulus modulation trajectory (Figs. 2C-D).

**Figure 2.**
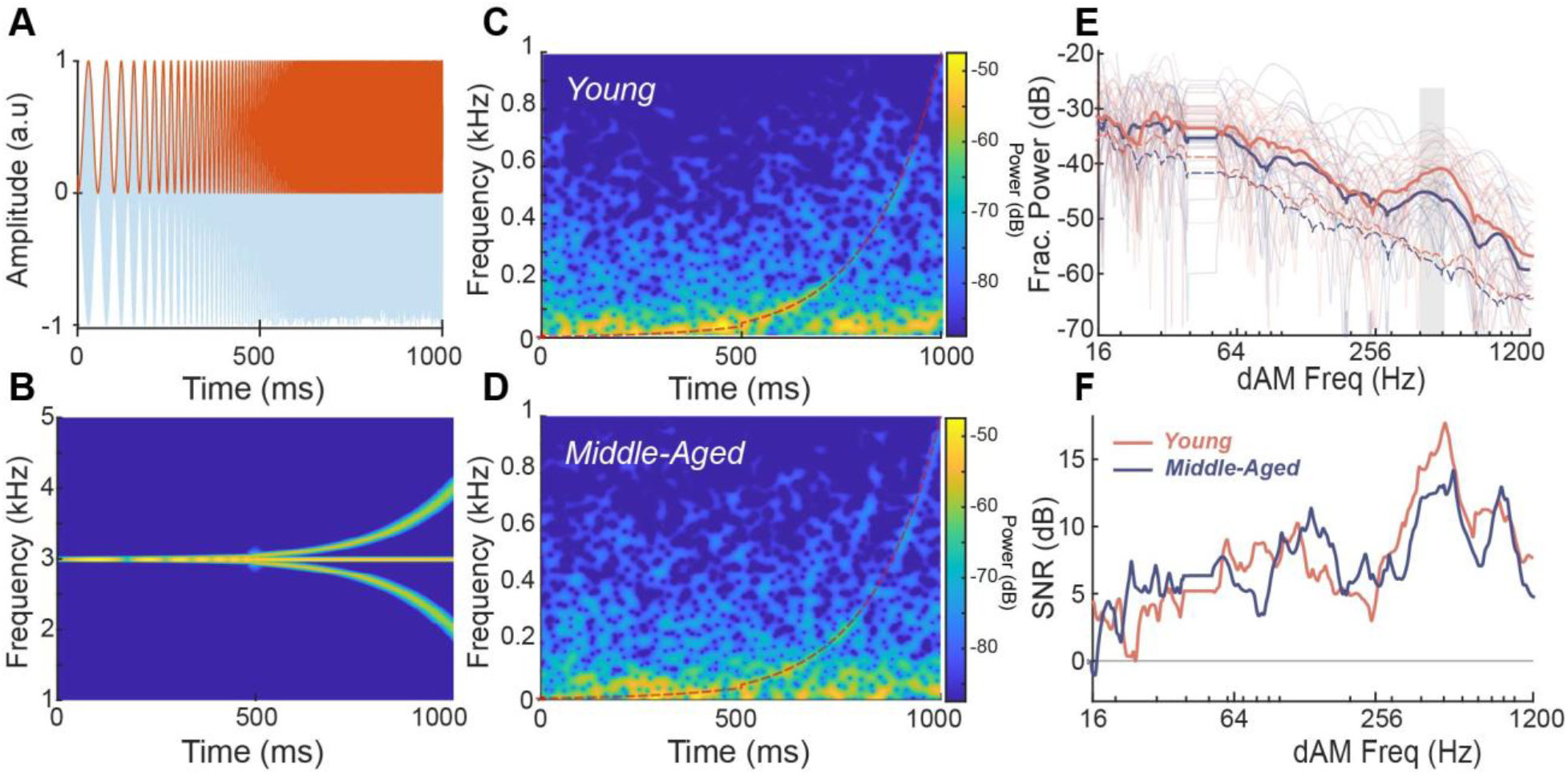
dAM EFRs provide a rapid, robust assessment of age-related temporal processing deficits. **(A)** Time-domain representation of the dAM stimulus. **(B)** Spectrogram of the dAM stimulus showing the exponentially sweeping modulation trajectory. **(C, D)** Representative dAM EFR response spectra in a young (C) and a middle-aged (D) gerbil, with the stimulus sweep overlaid (red). **(E)** Grand-average dAM EFR power along the modulation trajectory for young and middle-aged gerbils; middle-aged animals show reduced power at fast rates (gray bar). Light traces show individual animals. **(F)** dAM EFR signal-to-noise ratio versus modulation frequency, showing reliable responses at SNRs well above noise floor, and reduced SNRs in middle-aged animals at faster rates.

Spectrally specific frequency-demodulation analysis yielded robust dAM EFRs well above the noise floor across the modulation range in both groups (Figs. 2E-F). Young animals showed greater dAM EFR power than middle-aged animals, with the largest age-related differences at fast modulation rates above ∼400 Hz (Fig. 2E, gray bar). Signal-to-noise ratios confirmed reliable responses in both groups, with middle-aged animals showing reduced SNR at fast rates above 400 Hz (Fig. 2F). This age-related reduction at fast modulation rates closely mirrored the selective high-rate deficit observed with discrete sAM EFRs (Fig. 1E), validating dAM EFRs as an efficient measure of age-related temporal processing deficits.

### Machine-learning classification of dAM EFR features identifies the 400-500 Hz band as most discriminative

To objectively quantify the discriminability of dAM EFR features and identify the most informative spectral region, we applied a machine-learning classification framework (Fig. 3A). Spectral power was extracted along the dAM trajectory across the 20-1200 Hz modulation range and binned into contiguous frequency bands. A parametric search over candidate bin widths (20-400 Hz) across all the classifiers identified 100 Hz as optimal, yielding the highest classification performance across models (Fig. 3B). Using 100 Hz bins, nine classifiers were evaluated with leave-one-subject-out cross-validation, in which each animal is held out once for testing while the remaining animals are used for training. All classifiers performed above chance; XGBoost, a regularized gradient-boosting algorithm, achieved the highest mean *F*1score (*F*1 = 0.88), where *F*1 balances precision and recall (*34*) (Fig. 3C). Six out of the nine models consistently identified power in the 400-500 Hz band as the most informative feature for making the classification (Fig. 3D). SHAP (SHapley Additive exPlanations) analysis (*35*) of the best classifier showed the selective importance of power in the same 400-500 Hz band, with higher power associated with young animals and lower power with middle-aged animals (Fig. 3E). This band coincides with the largest age-related differences in the dAM grand averages (Fig. 2E) and with the higher rate sAM deficits (Fig. 1E), and it falls within the modulation range dominated by peripheral neural generators such as the auditory nerve. Critically, power in the 400-500 Hz band significantly correlated with 3 kHz synapse counts across animals *(Fig. 3F)*, relating the classifier’s key feature to histologically verified cochlear neural degeneration rather than an arbitrary statistical pattern. Overall, these results suggest that the dAM EFR classification reads out cochlear neural health directly rather than serving as a proxy for age *per se*.

**Figure 3.**
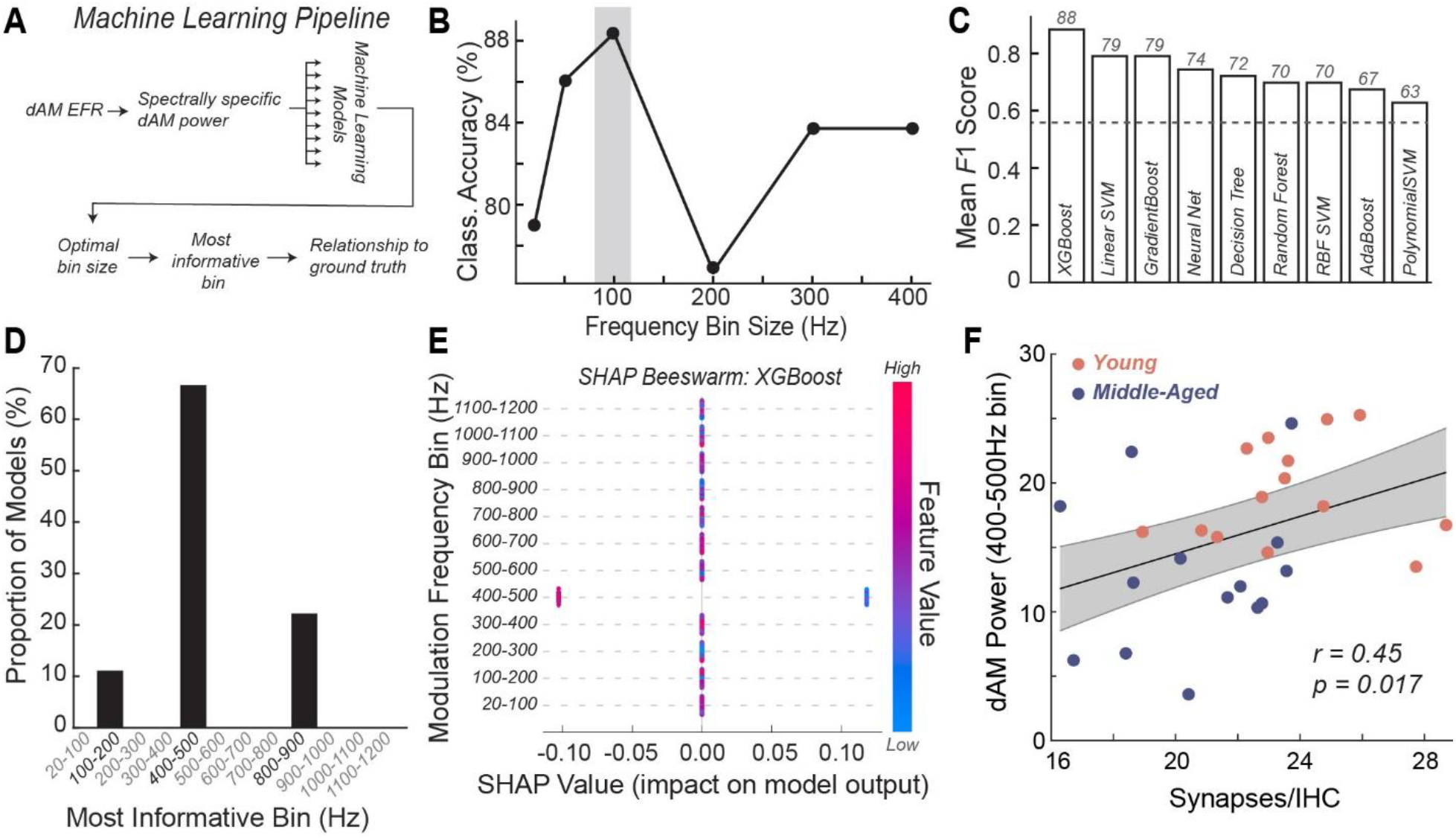
Machine-learning classification of dAM EFR spectral power identifies the 400-500 Hz band as the most informative feature distinguishing young from middle-aged gerbils. (A) Classification pipeline: spectrally specific dAM EFR power features are extracted across frequency bins and passed to machine-learning models to identify the optimal bin size and most discriminative band. (B) Classification performance across candidate bin sizes (20-400 Hz), for all models tested, peaking at 100 Hz (gray bar). (C) Mean *F*1 scores from leave-one-subject-out cross-validation across nine classifiers; XGBoost is highest (*F*1 = 0.88). (dashed line, 51% above chance). (D) Proportion of models identifying each frequency band as most informative; the 400-500 Hz band is selected by most models (E) SHAP beeswarm plot for the best-performing XGBoost classifier confirming the 400-500 Hz band as most discriminative; each point is one animal (color = relative feature value, red high / blue low). (F) Significant correlation between 3 kHz synapse counts and dAM EFR power in the 400-500 Hz band.

### Translational feasibility: the gerbil-derived signature transfers to human dAM EFRs

To assess the translational potential of the approach, we recorded dAM EFRs in 56 human listeners with clinically normal hearing: 25 young adults (18-25 years) and 31 middle-aged adults (40-55 years), using the same stimulus paradigm and spectrally specific analysis (Fig. 4A). Despite greater expected variability with human responses, middle-aged adults also showed reduced dAM EFR power across multiple regions including the 400-500 Hz modulation band relative to young adults (Fig. 4C), the same band identified in gerbils as the most discriminative classifier feature and the feature most strongly correlated with cochlear synapse counts. The SNR of the dAM EFR amplitude across modulation rates was lower than in gerbils, especially at rates greater than 500 Hz, yet predominantly well above the noise floor (Fig. 4D). We next tested whether the classification model itself transfers across species. The gerbil-trained classifier (Fig. 3) was applied directly to the human dAM EFR features without any retraining or human-specific adaptation. Despite being trained entirely on a different species, the model distinguished young from middle-aged adults above chance (Fig. 4E). These results establish proof-of-concept that the dAM EFR signature of CND is conserved across species and can be classified in humans using machine-learning approaches.

**Figure 4.**
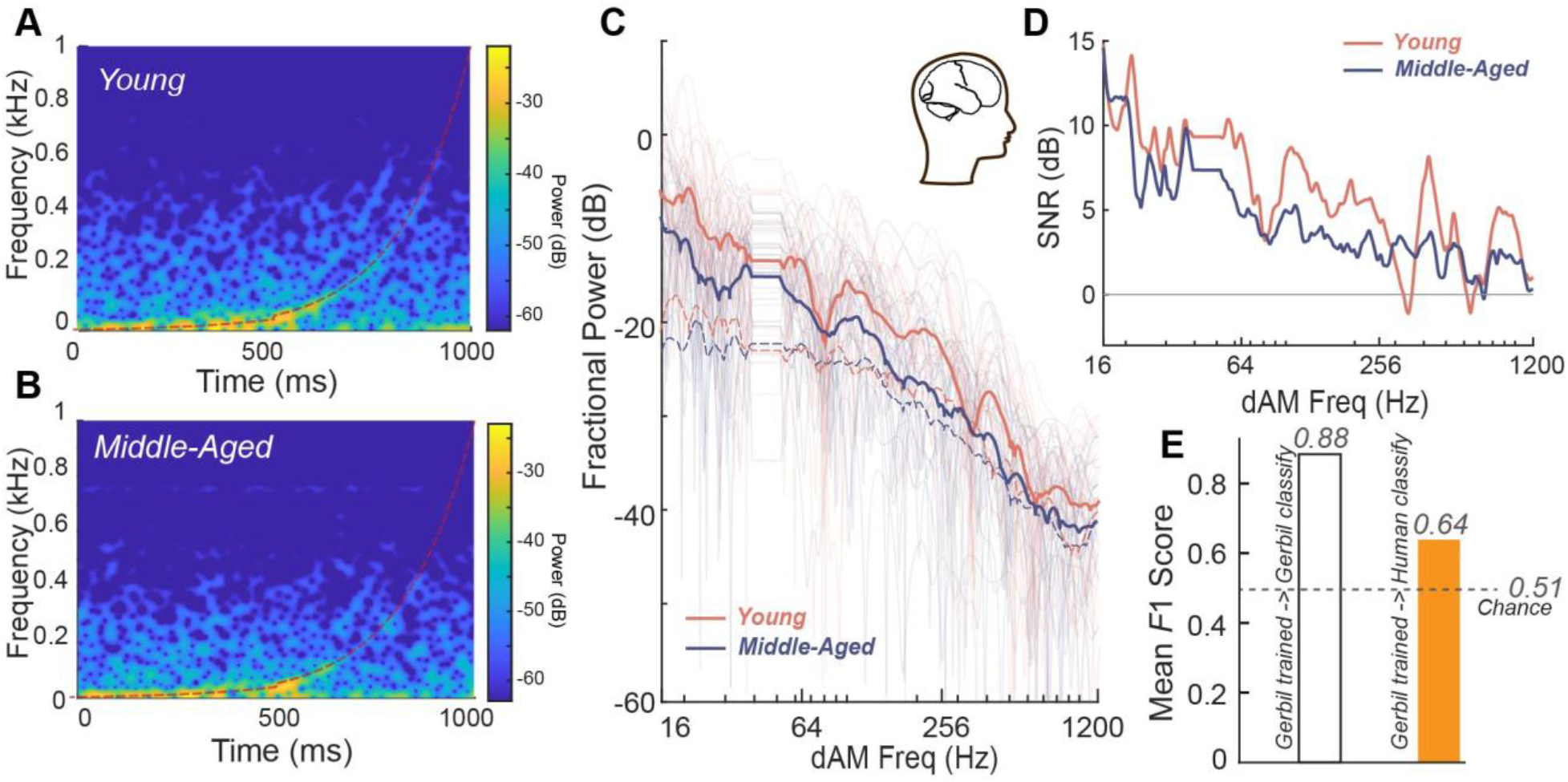
Translational feasibility of the dAM EFR CND signature in humans. (A-B) Grand-average dAM EFR power along the modulation trajectory in young (*n* = 25, A) and middle-aged (*n* = 31, B) adults with clinically normal audiograms. (C) Middle-aged adults showed reduced power in multiple bands including the 400-500 Hz band, mirroring the gerbil deficit (cf. Fig. 2E). (D) Response SNRs were above noise floor in humans. (E) Cross-species transfer with the gerbil-trained classifier (Fig. 3), applied to human dAM EFRs without retraining, distinguishes young from middle-aged adults above chance (dashed line, 51%, due to unequal sample sizes between the two groups).

## Discussion

We asked whether a novel dynamic amplitude-modulated envelope following response paradigm would be sensitive to age-related cochlear neural degeneration and whether the EFR spectral features can classify it. We found that (1) middle-aged gerbils exhibited cochlear synapse loss with preserved audiometric thresholds; (2) both discrete sAM and dAM EFRs captured selective amplitude reductions at fast modulation rates targeting putative auditory-nerve generators; and (3) machine-learning classification of dAM EFR spectral features discriminated age groups with high accuracy. The most discriminative feature, power in the 400-500 Hz band, correlated directly with histologically counted synapses. As an initial translational test, we applied the gerbil classifier directly to human data and achieved classification accuracy above chance.

Progressive loss of inner hair cell to auditory nerve fiber synapses is the earliest measurable form of CND, preceding hair cell loss and threshold elevation (*5, 10*), and is accompanied by selective EFR deficits at fast modulation rates (*18, 19, 29*). Our findings replicate this pattern in a species with low-frequency hearing that is well matched to humans, with histological confirmation and an expanded cohort relative to prior EFR aging studies in this model (*29*).

A central finding is the emergence of the 400-500 Hz dAM EFR power band as the most discriminative feature for classifying young from middle-aged animals, a result consistent across classifiers and corroborated by its correlation with synapse counts. Because phase-locking declines from periphery to cortex, fast-rate EFRs are dominated by auditory nerve and cochlear nucleus populations (*18, 19, 27*). Studies in mice established that EFRs generated at modulation frequencies spanning 400-1400 Hz are generated predominantly by these peripheral structures, as confirmed by group-delay analysis, and that deficits following neuropathic exposure are most prominent at these rates (*18, 19*). While broadly consistent with prior work, the specific frequency band diverges slightly from results obtained with discrete EFRs, which suggest that 1024 Hz AM rate is associated with the highest CND-related loss (*19, 29*). The dynamics of the different stimuli and the limited number of cycles of those faster modulation rates in the dAM stimulus are likely reasons for this difference. Nevertheless, the 400-500 Hz band sits at a mechanistically important transition, where it is fast enough that its generators are dominated by the peripheral neural generators most vulnerable to CND while still yielding sufficient signal-to-noise ratio in the dAM framework for stable individual-level estimates. Critically, the power in this band was correlated with histologically verified CND, suggesting that dAM EFRs are a promising biomarker to identify CND and strengthening the interpretation of the dAM EFR classifier as a readout for CND rather than just a proxy for age.

The dAM paradigm offers a route to clinical translation that conventional sAM approaches cannot readily provide. By sweeping continuously through modulation frequencies within a single brief stimulus, the dAM approach captures the full temporal modulation transfer function in a fraction of the time required by serial discrete measurements, while also providing substantially improved spectrotemporal resolution. Current diagnostic options for patients with normal audiograms and self-reported hearing difficulty are limited (*4, 36, 37*). Approximately 1 in 10 of these listeners have no objective clinical correlate of their complaint (*1*–*4*), and standard audiometry fails to capture the suprathreshold coding that supports everyday communication (*4, 38*–*40*). Combined with the machine-learning framework demonstrated here, dAM EFRs offer an objective, rapid, mechanistically grounded readout that could meaningfully expand the diagnostic toolkit. A natural next step is translation to humans, in whom dAM EFRs already yield robust, high-SNR responses.

Our human data provides initial evidence that this signature translates across species. Middleaged adults with normal audiograms showed the same 400-500 Hz dAM EFR reduction observed in gerbils, and a classifier trained solely on gerbil data classified human age groups above chance without any retraining. Given the substantial cross-species differences in head size, electrode montage, absolute response amplitude, and noise floor, the transfer of a gerbil-derived decision boundary to humans, even at modest accuracy, indicates that the underlying spectral signature of cochlear neural coding is conserved. Although this is a feasibility result rather than a diagnostic result, and given that human data carry no histological ground truth, the direction of the analysis is nonetheless informative. Because the gerbil classifier is anchored to ground truth synapse counts, its above-chance performance on humans reflects the presence of CND in the human response, whereas a classifier trained within the human cohort on age labels alone might instead exploit age-related features unrelated to CND. The classifier was also trained on the average response per animal, thus yielding a smaller sample size. Future studies will exploit the trial-by-trial variability of this response by training neural network-based classifiers on trial-level data. Additionally, converting this cross-species signal into an individual-level human diagnostic will require training directly on histologically verified CND across multiple etiologies and species and applying domain-adaptation transfer-learning methods that select features common to CND while suppressing species-specific variance, an approach that is also actively being pursued.

In summary, dAM EFRs detect age-related cochlear neural degeneration in Mongolian gerbils, recapitulating the selective high-rate deficits of conventional sAM EFRs in a fraction of the recording time. Machine-learning classification of dAM EFR spectral features discriminates age groups with high accuracy; the most discriminative feature, power in the 400-500 Hz band, is anchored to histologically verified synapse loss. Preliminary results also suggest feasibility of translation to humans. These results position the dAM EFR paradigm as a promising, mechanism-grounded foundation for rapid, objective clinical screening of hidden hearing disorders.

## Methods

### Experimental animals

Mongolian gerbil (*Meriones unguiculatus*) breeders were purchased from Charles River Laboratories and bred in-house. Twenty-four young Mongolian gerbils aged 16-46 weeks (male = 11, female = 13) and 19 middle-aged Mongolian gerbils aged 74-93 weeks (male = 6, female = 13) were studied. All animals were born and raised in the colony under climate-, temperature-, and light-controlled conditions. Data logging revealed an average ambient noise level of 56 dB SPL, with transients not exceeding 74 dB SPL during regular housing and up to 88 dB SPL during weekly cage changes. All procedures were approved by the Institutional Animal Care and Use Committee of the University of Pittsburgh (Protocol #24044677).

### Experimental setup

All procedures closely followed previously published studies (*27, 29, 41, 42*). Experiments were performed in a double-walled acoustic chamber. Animals were placed on a water-circulated warming blanket set to 37 °C with the pump outside the chamber to eliminate acoustic and electrical interference. Gerbils were initially anesthetized with isoflurane (4%) in an induction chamber, transferred and maintained at 1.5% isoflurane. Subdermal electrodes (Ambu) were placed with the positive electrode at the vertex, the negative electrode at the ipsilateral mastoid, and the ground at the base of the tail. After the positioning of the electrodes, animals were injected with dexmedetomidine (Dexdomitor, 0.3 mg/kg subdermally) and taken off isoflurane. Isoflurane exposure during setup was approximately 10 minutes; recordings commenced 15 minutes after cessation of isoflurane, with the washout period determined empirically (∼9 minutes) from ABR waveforms, latencies, and foot-pinch responses. Dexmedetomidine, an alpha-adrenergic agonist with sedative and analgesic effects, decreases motivation while preserving behavioral and neural responses, maintaining a compliant, unanesthetized-like state for ∼3 hours. Stimuli were presented to the right ear via insert earphones (ER3C, Etymotic). Electrode impedances were always <1 kΩ (RA4LI headstage, Tucker-Davis Technologies, TDT). Stimulus presentation and acquisition used a LabVIEW program. Output was calibrated using a Brüel & Kjær microphone and SigCal (TDT) to within ±3 dB across the frequency range tested. Waveforms were recorded with a multichannel system (RZ-5, TDT) and analyzed in BioSig or custom MATLAB programs (MathWorks).

### Stimulus presentation and recording

ABRs were elicited to tone bursts (0.5 ms rise/fall, 5 ms duration, 27.1/s, alternating polarity) at 0.5, 1, 3, and 8 kHz, and to a temporally compensated broadband chirp (a frequency sweep spanning 0.3-9 kHz in 2.1 ms) (*43, 44*), in which low frequencies are delivered earlier than high frequencies to offset cochlear traveling-wave delay, producing synchronous stimulation across the cochlea and larger, more easily identified waveforms. Sound level was increased from 20 to 80 dB SPL in 10 dB steps. Responses were amplified 10,000× amplification; TDT Medusa 4z) and filtered (0.3-3 kHz). Trials exceeding 15 µV were rejected; 512 artifact-free trials of each polarity were averaged. ABR thresholds were defined as the minimum level producing a visually detectable response, assessed by two blinded, trained observers.

Discrete EFRs were elicited to sinusoidally amplitude-modulated (sAM) tones (5 ms rise/fall, 250 ms duration, 3.1/s, alternating polarity) at a 3 kHz carrier. Modulation frequency varied from 16 to 1024 Hz at a fixed level of 30 dB SL (above individual 3 kHz ABR threshold), and in a separate experiment the level was varied from 20 to 80 dB SPL at a fixed 1024 Hz modulation frequency. Responses were amplified (10,000× amplification; TDT Medusa 4z) and filtered (0.1–10 kHz). Trials exceeding 15 µV were rejected; 250 artifact-free trials of each polarity were averaged. Fast Fourier Transforms (FFTs) were computed on average waveforms beginning 10 ms after onset (to exclude the ABR) through offset. Peak FFT amplitude was taken as the maximum of three 3 Hz bins around the modulation frequency.

dAM EFRs were elicited to dynamically amplitude-modulated stimuli with a 3 kHz carrier. For a desired modulation-frequency trajectory *f*_*am*_*(t)*, the phase trajectory φ_am_(*t*) is its time integral, and the stimulus is,

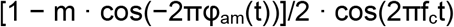

with carrier frequency f_c_ and modulation depth m. The stimulus comprised two 500 ms segments: in the first, *f*_*m*_ increased exponentially from 2.5 to 40 Hz (range with substantial cortical contributions); in the second, from 50 to 1440 Hz (predominantly subcortical contributions). The exponential sweep approximates the logarithmic modulation processing of the auditory system. dAM stimuli were presented at 0.8/s, with approximately 200 sweeps collected per animal.

dAM EFRs were analyzed using spectrally specific frequency-demodulation methods that estimate power along a known time-varying trajectory. For a signal x(t) and trajectory *f*_*am*_*(t)*, the analytic signal was obtained via the Hilbert transform; *a(t)* was defined as 1/√2 times its pointwise magnitude. The trajectory phase

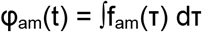

was used to frequency-demodulate a(t) so that power along *f*_*am*_*(t)* maps to 0 Hz. Power was estimated as the low-frequency power of the demodulated signal using a multitaper spectrum (MATLAB pmtm; time-bandwidth half-product = 1.5) within a 24 Hz band (−12 to 12 Hz). A noise floor was estimated from a parallel trajectory shifted up by 36 Hz to reduce overlap.

### Immunohistology

Procedures closely followed previously published studies (*19, 29*). Deeply anesthetized gerbils were transcardially perfused with 4% paraformaldehyde in 0.1 M phosphate buffer, followed by intralabyrinthine perfusion of both cochleas. Cochleas were post-fixed (4% paraformaldehyde, 1 h), decalcified in 0.12 M EDTA (5-7 days), microdissected into half-turns, and immunostained for C-terminal binding protein 2 (mouse anti-CtBP2, BD Biosciences, 1:200), glutamate receptor 2 (mouse anti-GluR2, Invitrogen, 1:2000), and myosin-VIIa (rabbit anti-myosin-VIIa, Proteus Biosciences, 1:200), with Alexa Fluor conjugated secondary antibodies.

Cochlear pieces were measured and a frequency map was computed using a custom ImageJ plug-in. Confocal z-stacks at the 0.5, 1, 3, and 8 kHz regions were collected on a Nikon A1 microscope (z-stacks spanning the cuticular plate to the synapse of ∼10 hair cells in each frequency, 0.25 µm z-steps, 1024 × 1024 raster, 60× oil-immersion lens objective, 1.59× digital zoom). Stacks were analyzed in Imaris (Oxford Instruments); inner hair cells were identified by myosin-VIIa and presynaptic ribbons and postsynaptic patches were counted from 3D reconstructions. Synaptic ribbons per inner hair cell in each targeted frequency region was calculated.

### Human participants and dAM EFR recordings

Human dAM EFRs were recorded in 56 adults with clinically normal audiograms: 25 young adults (18-25 years) and 31 middle-aged adults (40-55 years). Young and middle-aged participants were recruited and screened using IRB approved protocols (University of Pittsburgh Institutional Review Board #21040125). All participants provided informed consent and were compensated. Recruitment drew on the University of Pittsburgh Pitt+Me research registry, the University of Pittsburgh Department of Communication Science and Disorders participant pool, and the broader Pittsburgh community. Eligibility was determined at an initial session and required normal cognition (*Montreal Cognitive Assessmen*t ≥25), normal-hearing thresholds (≤25 dB HL, 250-8000 Hz), no severe tinnitus (*Tinnitus Handicap Inventory*), loudness discomfort levels > 80 dB HL at 0.5, 1, and 3 kHz, and self-reported American English fluency. Both age groups underwent identical audiological characterization, including otoscopy, standard and extended high-frequency audiometry (up to 16 kHz), loudness discomfort levels, and a self-reported noise-exposure history, as reported in a previously published study from our group (*29*).

Recordings were made in a sound-attenuating booth with a 64-channel EEG system (BioSemi ActiveTwo) while participants reclined and watched a silent, subtitled film. dAM stimuli (3 kHz carrier, sweeping the modulation trajectory described above) were generated with an RX6 processor (TDT) and delivered at 85 dB SPL through ER3C insert earphones (Etymotic) coupled to gold-foil tiptrodes seated in the ear canals, which additionally recorded ear-canal potentials. External triggers synchronized the stimulus and recorded EEG for offline epoching. Consistent with the montage used to establish the dAM paradigm in humans, EFRs were analyzed from the vertex electrode (Fz) referenced to the ipsilateral tiptrode using custom MATLAB scripts, and each participant contributed a single recording. dAM EFR power was estimated with the same spectrally specific frequency-demodulation analysis applied to the gerbil data.

### Machine-learning classification

Spectral power features were extracted from dAM EFRs by computing power along the dAM trajectory within contiguous frequency bands spanning the 20 to1200 Hz modulation range. A data-driven search over candidate bin sizes (20, 50, 100, 200, 300, and 400 Hz) selected the size producing the highest classifier F1-score across models (100 Hz). Nine classifiers were trained: Linear SVM, RBF SVM, Polynomial SVM, Decision Tree, Random Forest, Gradient Boosting, AdaBoost, XGBoost, and Multi-Layer Perceptron. A leave-one-subject-out cross-validation scheme held out one animal at a time for testing. Hyperparameters were optimized by grid search using the highest mean *F*1score across folds, where

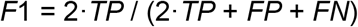

Where TP is True Positive, FP is False Positive and FN is False Negative. Feature importance was assessed with SHAP values (SHapley Additive exPlanations)(*35*), where larger absolute values indicate greater contribution and the sign indicates the favored class; beeswarm plots visualize the direction and magnitude of each feature’s contribution across animals, color-coded by feature value.

For the cross-species analysis (Fig. 4), human dAM EFRs were processed through the identical feature pipeline used for gerbils, namely spectrally specific power along the dAM trajectory binned into the same 100 Hz frequency bands, to yield a feature vector matched in dimension and definition to the gerbil training features. The gerbil-trained XGBoost classifier (Fig. 3) was then applied to these human features without any retraining, fine-tuning, or human-specific adaptation. To place gerbil and human features on a common scale despite species differences in absolute response amplitude, electrode montage, and noise floor, features were normalized within species prior to classification. Classification accuracy for distinguishing young from middle-aged adults was evaluated against the chance level using a label-permutation test with a 95% confidence interval. Chance level was estimated by randomly permuting the class labels 10,000 times and computing the mean accuracy against the true labels.

### Statistical analysis

The study was cross-sectional. Statistical outliers were identified and removed using Tukey’s fence rule (k = 1.5). Descriptive statistics were computed in Microsoft Excel, inferential analyses used the Python *pingouin* library, and correlations were computed in MATLAB (Pearson’s *r*, with degrees of freedom, r, and p reported). EFR amplitude was analyzed across modulation frequency (16, 40, 110, 256, 512, and 1024 Hz) and sound level (20-80 dB SPL), comparing young and middle-aged animals. Normality was assessed with the Shapiro-Wilk test (α = 0.05) and homogeneity of variance with Levene’s test. Two-way mixed ANOVAs used age group as the between-subjects factor and modulation frequency or sound level as the within-subjects factor, with Greenhouse-Geisser correction for sphericity violations. Where normality was violated, Mann-Whitney U tests were used for follow-up comparisons with Bonferroni correction across conditions. Partial eta-squared (η_p_^2^) is reported as effect size.

## Author contributions

A.P. conceived and supervised the study and acquired funding. L.S.A. performed the gerbil electrophysiology experiments and analyzed the data. R.G. developed and performed the machine-learning classification analyses. J.K. performed the gerbil immunohistology. M.A. supervised the machine-learning and signal-processing methodology. E.L.B. contributed to the dynamic amplitude-modulated stimulus methods and analysis pipelines. A.P. and L.S.A. wrote the manuscript with input from all authors. All authors reviewed and approved the final manuscript.

## Acknowledgements

This work was supported by the National Institute on Deafness and Other Communication Disorders, National Institutes of Health (R21DC018882 and R01DC023721, PI: Parthasarathy; T32NS126122, trainee: Andrino) and the Department of Defense (HRRP W81XWH2110602, PI: Bartlett). We thank Dr. Simon Watkins and the Center for Biological Imaging at the University of Pittsburgh for confocal imaging support (S10RR028478), CTSI for recruitment support (UL1TR001857) and Maggie Zink, Jacie McHaney, Rachel Blake, Caroline Henning, Kimberly Yurasits, Claire Mitchell, and Victoria Cancel for assistance with data collection. The dynamic AM stimuli and spectrally specific analysis methods are patented but available for non-commercial use (PCT International Application No. PCT/US2024/011312, claiming priority from U.S. Provisional Application No. 63/479,768).

## Data and code availability

Data and code used in this manuscript are available for non-commercial use will be available here upon publication - https://osf.io/dwr9b/

